# Aminoglycoside heteroresistance in *Enterobacter cloacae* is driven by the cell envelope stress response

**DOI:** 10.1101/2023.10.11.561937

**Authors:** Ana J. Choi, Daniel J. Bennison, Esha Kulkarni, Haoyu Sun, Hanqi Li, Jonathan Bradshaw, Hui Wen Yeap, Nicholas Lim, Vishwas Mishra, Anna Crespo-Puig, Frances Davies, Shiranee Sriskandan, Avinash R. Shenoy

**Author notes:** equal contribution (Ana J. Choi, Daniel J. Bennison). Correspondence & Lead Contact: Address: Room 4.40A, Flowers Bldg, Armstrong Road, MRC CMBI, Imperial College London, London SW7 2AZ, UK.

## Abstract

*Enterobacter cloacae* is a Gram-negative nosocomial pathogen of the ESKAPE priority group with increasing multi-drug resistance via the acquisition of resistance plasmids. However, *E. cloacae* can also display phenotypic antimicrobial resistance, such as heteroresistance or persistence. Here we report that *E. cloacae* ATCC 13047 and six strains isolated from patients with blood infections display heteroresistance or persistence to aminoglycosides. *E. cloacae* heteroresistance is transient, accompanied with formation of ‘petite’ colonies and increased MIC against gentamicin and other aminoglycosides used in the clinic, but not other antibiotic classes. To explore the underlying mechanisms, we performed RNA sequencing of heteroresistant bacteria, which revealed global gene-expression changes and a signature of the CpxRA cell envelope stress response. Deletion of the *cpxRA* two-component system abrogated aminoglycoside heteroresistance and petite colony formation, pointing to its indispensable role in phenotypic resistance. The introduction of a constitutively active allele of *cpxA* led to high aminoglycoside MICs, consistent with cell envelope stress driving these behaviours in *E. cloacae*. Cell envelope stress can be caused by environmental cues, including heavy metals. Indeed, bacterial exposure to copper increased gentamicin MIC in the wild type, but not the Δ*cpxRA* mutant. Moreover, copper exposure also elevated the gentamicin MICs of bloodstream isolates, suggesting that CpxRA- and copper-dependent aminoglycoside resistance is broadly conserved in *E. cloacae* strains. Altogether, we establish that *E. cloacae* relies on transcriptional reprogramming via the envelope stress response pathway for transient resistance to a major class of frontline antibiotic.

**Importance:** *Enterobacter cloacae* is a bacterium that belongs to the ESKAPE priority group and an increasing threat worldwide due its multidrug resistance. *E. cloacae* can also display phenotypic resistance to antibiotics, leading to treatment failure. We report that sensitive strains of *E. cloacae,* including six strains isolated from patients with bloodstream infections, show heteroresistance or persistence to aminoglycoside antibiotics. These are important frontline microbicidal drugs used against Gram-negative bacterial infections, therefore understanding how resistance develops in sensitive strains is important. We show that aminoglycoside resistance is driven by the activation of the cell envelope stress response and transcriptional reprogramming via the CpxRA two-component system. Further, heterologous activation of envelope stress via copper, typically a heavy metal with antimicrobial actions, also increased aminoglycoside MICs in all tested strains of *E. cloacae*. Our study suggests phenotypic aminoglycoside resistance in *E. cloacae* could be broadly conserved and cautions against the undesirable effects of copper.

## Introduction

*Enterobacter cloacae* is a Gram-negative opportunistic pathogen that can cause life-threatening hospital-acquired infections (1, 2). These include bloodstream infections and sepsis, peritonitis, lower respiratory tract and urinary tract infections. Due to its natural and acquired resistance to several classes of antibiotic, *E. cloacae* is included in the ESKAPE (*Enterococcus, Staphylococcus, Klebsiella, Acinetobacter, Pseudomonas and Enterobacter spp*) group of priority multidrug-resistant pathogens (1, 3). *E. cloacae* is the type species of *Enterobacter* and the third leading cause of death from drug-resistant *Enterobacteriaceae* infection after *Escherichia coli* and *Klebsiella pneumoniae* worldwide (4), and often present as nosocomial bloodstream infections in both adults and neonates (5, 6). Therefore, a better understanding of mechanisms of antimicrobial resistance in *E. cloacae* is critical for ensuring continued efficacy of current frontline drugs.

Antibiotic resistance typically evolves through acquisition of plasmids encoding antibiotic-inactivating enzymes, mutation of antibiotic targets, increased drug efflux, and limiting drug uptake (7). These mechanisms result in a stable, heritable increase in the minimum inhibitor concentration (MIC) of an antibiotic, which subsequently becomes ineffective in treatment. Less common, but equally formidable, threats to treatment are heteroresistance and persistence in susceptible strains that transiently become drug-resistant (8, 9). These behaviours emerge from heterogeneity in clonal bacterial cultures wherein most of the population is susceptible and killed by an antibiotic, but a subpopulation survives antibiotic exposure. Such population heterogeneity results in a characteristic biphasic antibiotic time-to-kill curve, with rapid early killing of the majority and slower killing of the resistant subpopulation (8). Heteroresistance and persistence differ in that heteroresister, but not persister, bacteria display high MIC against the antibiotic and can grow in its presence (8). Both mechanisms are typically ‘unstable’ as the surviving bacteria produce heterogenous populations containing susceptible and resistant bacteria, which indicates that these forms of resistance are also heritable. Phenotypic resistance may facilitate the acquisition of genetically stable antibiotic resistance, and as such further understanding of the underlying mechanisms behind this resistance is critical to improving the efficacy of treatment (10).

Increasing antibiotic resistance is a major concern in *E. cloacae*, which is naturally resistant to most β-lactams (including first, second and third-generation cephalosporins) via the AmpC cephalosporinase (11), and is fast acquiring carbapenem-resistance plasmids (12). Plasmid-acquired resistance to other antibiotics, including polymyxins and fosfomycin, is also on the rise (13, 14). Molecular mechanisms leading to heteroresistance remain poorly understood, however, unstable increase in the copy number of antibiotic-resistance or - modifying genes have been reported (8, 15). Non-genetic mechanisms are also recognised, including increased expression of lipopolysaccharide (LPS)-modifying enzymes via the PhoPQ two-component system leading to colistin heteroresistance (13). However, mechanisms of heteroresistance against aminoglycosides are not known.

Aminoglycosides bind to the 30S ribosomal subunit and interfere with protein translation. However, unlike other drugs that target the ribosome, aminoglycosides are bactericidal, probably due to error-prone translation and protein misfolding which lead to toxicity (16). As aminoglycosides are charged molecules that do not easily enter eukaryotic cells, they are often used in ‘gentamicin-protection assays’ to kill extracellular bacteria and measure bacteria that invaded into host cells and are protected from the antibiotic (17–19). During our studies on macrophage infection by the gentamicin-sensitive type-strain of *E. cloacae*, namely ATCC 13047 (originally isolated from a spinal fluid specimen), we were surprised that extracellular bacteria were not eliminated by 100 mg.L^−1^ gentamicin even at 5 h post-treatment. As aminoglycosides are integral to treatment options against sepsis and multidrug-resistant Gram-negative infections (20), we decided to investigate this unexpected behaviour further. Here we report aminoglycoside heteroresistance in the *E. cloacae* type-strain, ATCC 13047 (21) and persistence in aminoglycoside-sensitive isolates collected from adult bloodstream infections as part of the Bioresource in Adult Infectious Diseases (BioAID) cohort study (22). Heteroresistance is associated with a subpopulation that forms unstable petite colonies that we term gentamicin-associated petite *E. cloacae* (GAPEc). We find that increased expression of the cell envelope stress response by the CpxRA two-component system is indispensable for heteroresistance. We further show that activation of CpxRA by copper also triggers aminoglycoside resistance in *E. cloacae* clinical isolates. Our studies highlight potential risks and provide clarity on mechanisms of phenotypic aminoglycoside resistance in *E. cloacae*.

## Results

### A subpopulation in clonal E. cloacae cultures displays gentamicin resistance

We initially observed incomplete killing of extracellular *E. cloacae* ATCC 13047 during gentamicin-protection assays in cell culture experiments, and proceeded to confirm that this strain is susceptible to gentamicin and does not grow when inoculated in LB containing 10-20 mg.L^−1^ gentamicin (∼10-20x MIC). However, to our surprise, in disc-diffusion assays alongside a gentamicin-susceptible reference strain of *E. coli* (ATCC 11775) we observed colonies of *E. cloacae* within the zone of clearance (***Figure 1A-B***). As expected, *E. coli* showed zones of clearance ≥ 17 mm in diameter, indicating susceptibility to gentamicin based on EUCAST guidelines (23). The formation of an ‘inner’ zone ≤ 17 mm suggested that *E. cloacae* displays a form of phenotypic resistance to gentamicin. Notably, exponential and stationary phase *E. cloacae* showed similar behaviour in disc-diffusion assay (***Figure 1B***), indicating that observed resistance in disc-diffusion assays did not depend on the stage of growth. As these observations were suggestive of a heterogenous population of sensitive and resistant subpopulations, we investigated this phenomenon further using time-to-kill and broth microdilution-based MIC assays which are more reliable in detecting persister or heteroresister subpopulations.

**Figure 1.**
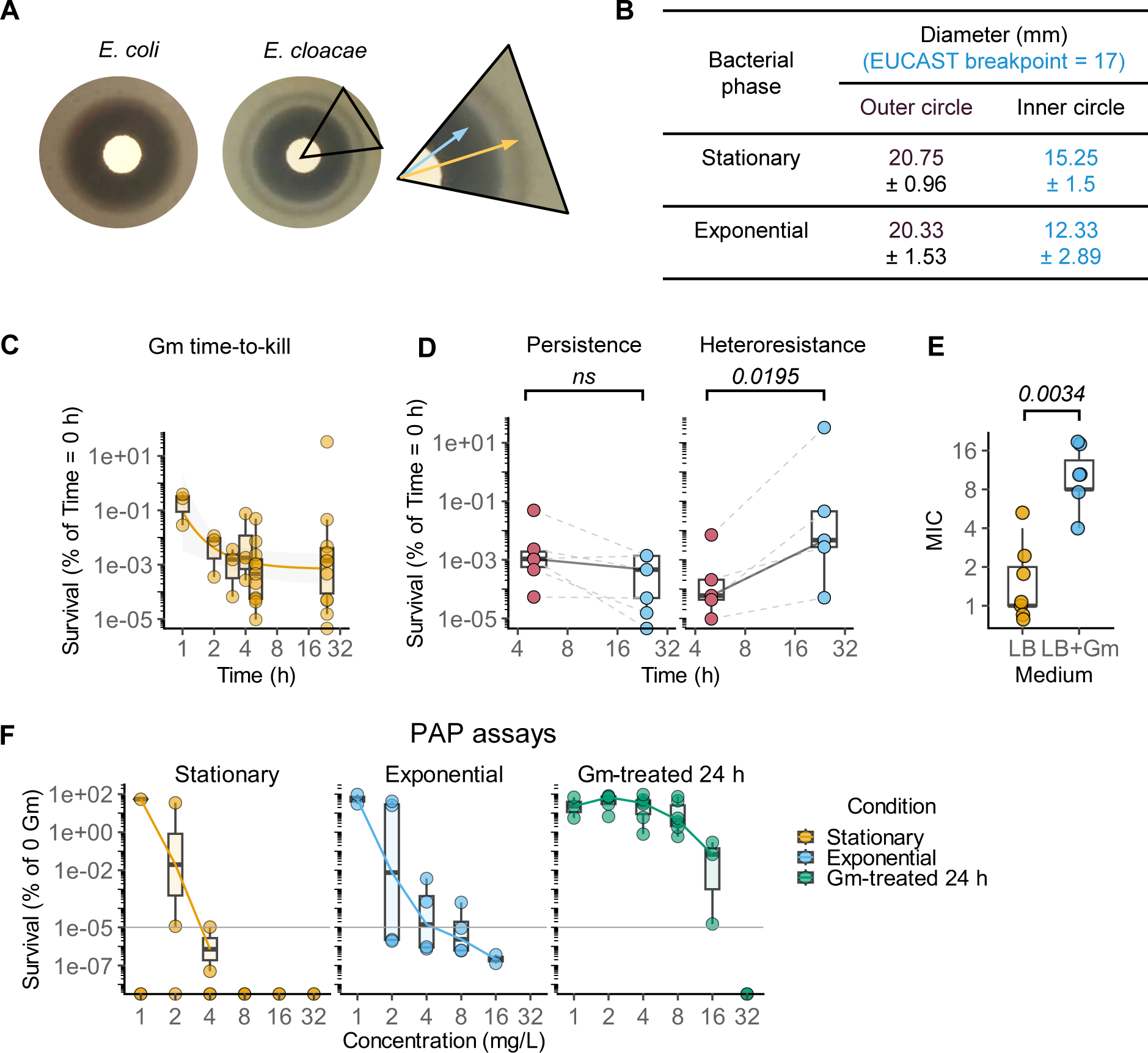
*E. cloacae* contains a subpopulation that is resistant to gentamicin. **(A-B)** Representative images from disc-diffusion assays testing gentamicin sensitivity of *E. coli* ATCC 11775 and *E. cloacae* ATCC 13047 (**A**). Inset shows *E. cloacae* colonies inside the zone of clearance, whose diameters are shown in (**B**). Mean ± SD of independent experiments indicated; n = 4 for stationary phase and n = 3 for exponential phase cultures. **(C-D)** Gentamicin (Gm) time-to-kill curves showing the percentage survival of *E. cloacae* over time. Cultures were treated with gentamicin (20 mg.L^−1^), samples collected, washed and plated on LB agar to quantify viable CFU.mL^−1^. Graph in (**C**) shows first order decay curve fit to data. Graphs in (**D**) show percentage of viable bacteria at 5 and 24 h post-gentamicin exposure showing persistence-like and heteroresistance-like behaviour as labelled. Box and whiskers with median are plotted; number of independent experiments as follows: n = 11 in **C**; n = 5-6 each in **D**. Two-tailed *P* value from mixed effects ANOVA; ns = not significant. **(E)** MIC measured by broth microdilution for bacteria grown in lysogeny broth (LB) without or with gentamicin (20 mg.L^−1^) for 24 h (Gm). Box and whiskers with median are plotted from n = 7 (LB) and n = 6 (Gm) independent measurements. Two-tailed *P* value from a Mann-Whitney test. **(F)** Population analysis profiling (PAP) assays of *E. cloacae* ATCC 13047 cultures in stationary or exponential phase, or following gentamicin treatment (20 mg.L^−1^) for 24 h, as labelled. Bacteria were plated on Mueller Hinton broth (MHB) agar plates with the indicated concentrations of gentamicin. Percentage survival relative to plates without antibiotic are plotted. Box and whiskers with median from n = 4 independent experiments shown.

Treatment of exponentially growing *E. cloacae* with gentamicin led to ∼10^4^ – 10^5^-fold reduction in viable bacterial colony-forming units (CFU) between 2-5 h post-treatment (***Figure 1C***). Analyses of data from time-to-kill assays indicated biphasic killing-kinetics by gentamicin, with rapid early killing and a slower late phase (***Figure 1C***). Notably, across independent experiments, survival at 5 h post-treatment was 0.006 ± 0.015 % which increased to 3 ± 10 % at 24 h post-treatment (***Figure 1C***), pointing to an increase in the proportion of surviving bacteria between 5 and 24 h. We also noticed the variability between biological replicates and inspected these data closer, which revealed two trends. Survival between 5 and 24 h either remained unchanged (n = 6 experiments, ***Figure 1D***) indicating persistence-like behaviour, or increased (*P =* 0.0195; n = 5 experiments, ***Figure 1D***), indicating heteroresistance-like behaviour. Based on these experiments, we conclude that a subpopulation of susceptible *E. cloacae* displays antibiotic resistance that manifests as common forms of phenotypic resistance.

To distinguish between heteroresistance and persistence, we measured the MIC of gentamicin for bacteria that had survived exposure for 24 h. The median MIC of *E. cloacae* was ∼1 mg.L^−1^ (expectedly, lower than EUCAST breakpoint of 2 mg.L^−1^), which increased to ∼8 mg.L^−1^ (higher than EUCAST breakpoint; ***Figure 1E***) for bacteria that survived exposure to gentamicin. These results indicating an increase in MIC are consistent with the presence of a heteroresistant subpopulation in the culture.

Next, we independently verified heteroresistance using population analysis profiling (PAP) assays. PAP assays are designed to detect whether a bacterial population contains a subpopulation at a frequency of ≥ 10^−5^ % that can survive antibiotic at 8-fold higher concentration than the MIC (8, 24). Indeed, we observed ≥10^−5^ % bacteria could produce colonies on MHB plates containing up to 8 mg.L^−1^ gentamicin, confirming heteroresistance in *E. cloacae* (***Figure 1F***). Surprisingly, heteroresistance was only observed in exponentially growing cultures and not stationary-phase cultures (***Figure 1F***). Furthermore, in cultures exposed to 20 mg.L^−1^ gentamicin for 24 h (as in ***Figure 1B***), ∼0.1 % bacteria produced colonies on MHB agar containing 16 mg.L^−1^ gentamicin (***Figure 1F***), which further points to the proliferation and enrichment of the resistant subpopulation over time. Altogether, we conclude that exponentially growing *E. cloacae* contain a subpopulation with a higher MIC that withstands prolonged exposure to gentamicin concentrations that are typically bactericidal.

### Clinical E. cloacae strains display gentamicin persistence

We wanted to rule out that this behaviour could be limited to a laboratory strain of *E. cloacae* and therefore tested strains isolated from bloodstream infections. We used 7 strains isolated from patients in London as part of BioAID between 2015-2019 (***Table 1***); as ECL_AS01 was gentamicin-resistant, we did not include this strain in further studies. We assessed whether the six gentamicin-sensitive strains (ECL_AS02-ECL_AS07) showed phenotypic resistance to gentamicin. Disc-diffusion assays confirmed the six strains we tested were sensitive to gentamicin (***Figure 2A***); however, strain ECL_AS04 showed a large variation in the zone of clearance. None formed colonies inside the zone of clearance. We then assessed whether these strains can withstand 20 mg.L^−1^ gentamicin exposure for 20 h. Interestingly, all six strains showed a similar fraction of viable population at 5 and 20 h post-gentamicin treatment (∼10^−3^-10^−4^ % survival; ***Figure 2B***), which is similar to the persister-like behaviour observed with the ATCC 13047 strain (***Figure 1C-D***). From these experiments we conclude that gentamicin-sensitive clinical strains of *E. cloacae* can contain persisters that can survive exposure to ∼20x MIC.

**Figure 2.**
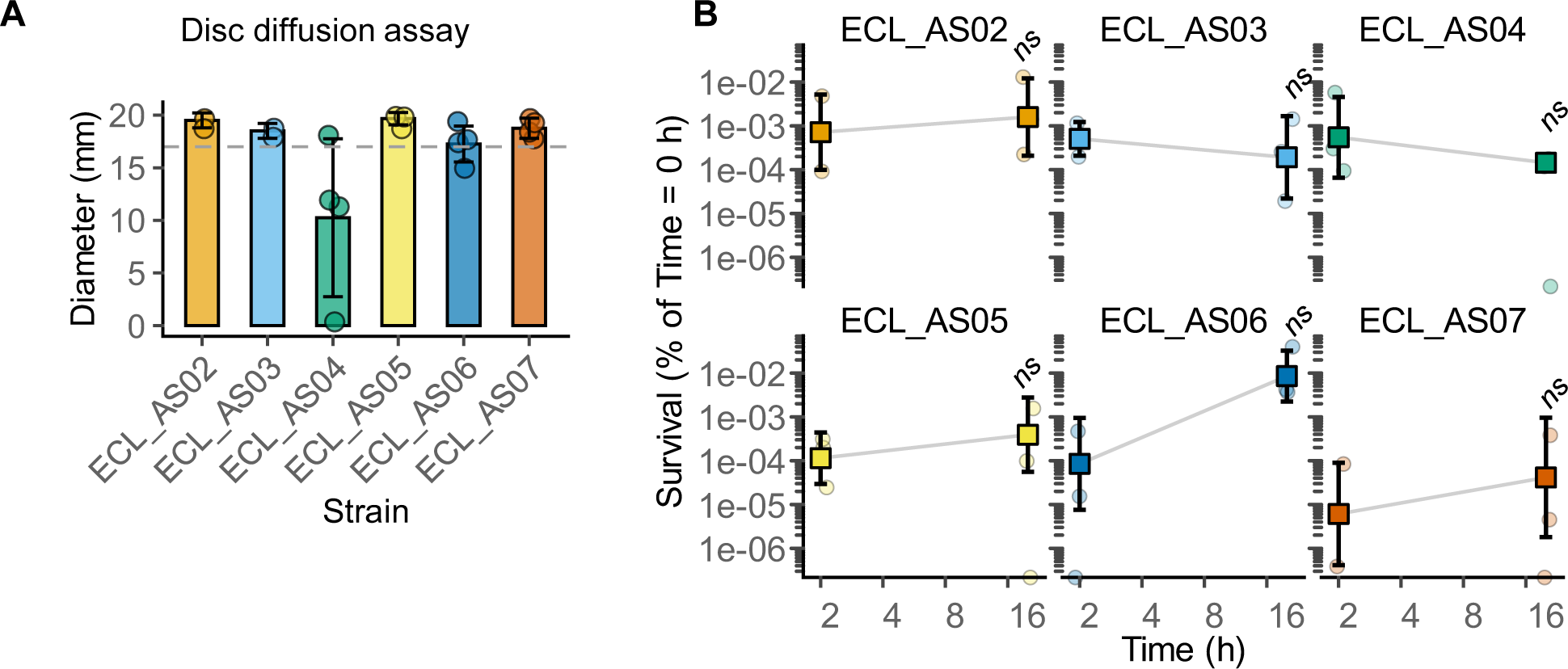
Clinical strains of *E. cloacae* display persistence against gentamicin. **(A)** Diameters of zone of clearance from gentamicin disc-diffusion assays for the indicated strains of *E. cloacae*. Mean and SD shown from n = 3-4 independent experiments. **(B)** Survival of the indicated clinical isolates of *E. cloacae* at the indicated time points after exposure to gentamicin. Exponentially growing cultures were treated with gentamicin (20 mg.L^−1^), washed twice and plated on LB agar to enumerate surviving bacteria. Mean and SD from n = 3 experiments are shown. ns, not significant, i.e., two-tailed *P* > 0.05 for the comparison between the two time-points for each strain by mixed effects ANOVA.

**Table 1.** Details and antibiotic susceptibility of clinical strains from the BioAID cohort.

| Strain ID | ECL_AS01 | ECL_AS02 | ECL_AS03 | ECL_AS04 | ECL_AS05 | ECL_AS06 | ECL_AS07 |
| --- | --- | --- | --- | --- | --- | --- | --- |
| Pathogen Isolated | *E. cloacae* | *E. cloacae* | *E. cloacae* | *E. cloacae* | *E. cloacae* | *E. cloacae* | *E. cloacae* |
| Date Collected | 04/2015 | 07/2015 | 05/2016 | 03/2017 | 03/2017 | 11/2017 | 09/2019 |
| ESBL Markers | Y | ND | N | ND | ND | N | ND |
| AmpC Markers | ND | ND | Y | ND | ND | Y | ND |
| Gentamicin | R | S | S | S | S | S | S |
| Amikacin | S | S | S | S | S | S | S |
| Tobramycin | R | S | S | S | S | S | ND |
| Amoxycillin | R | R | R | R | R | R | R |
| Augmentin (Co-amoxiclav) | R | R | R | R | R | R | R |
| Aztreonam | R | S | S | S | S | S | S |
| Cefotaxime/Ceftriaxone | R | S | S | S | S | S | S |
| Cefoxitin | R | S | R | S | R | R | R |
| Ceftazadime | R | S | S | S | S | S | S |
| Ceftazidime-avibactam | ND | ND | ND | ND | ND | ND | S |
| Cefuroxime | R | S | S | S | S | S | S |
| Ciprofloxacin | R | S | S | S | S | S | S |
| Colistin | S | S | S | ND | ND | S | ND |
| Cotrimoxazole | ND | ND | ND | ND | ND | ND | S |
| Ertapenem | S | S | S | S | S | S | S |
| Meropenem | S | S | S | S | S | S | S |
| Tazocin (piperacillin tazobactam) | R | S | S | S | S | S | S |
| Temocillin | S | S | S | S | S | S | S |
| Trimethoprim | R | S | S | S | S | S | ND |
| Tigecycline | S | S | S | S | S | S | ND |

S = sensitive; R = resistant, ND = not determined

### Gentamicin-associated petite E. cloacae (GAPEc) colony phenotype confers heteroresistance

In gentamicin time-to-kill experiments (***Figure 1C***) we noticed the appearance of a mix of ‘petite’ and ‘normal’ sized colonies of *E. cloacae* upon plating on antibiotic-free LB agar plates (***Figure 3A-B***). Here on, we call the smaller colonies gentamicin-associated petite *E. cloacae* (GAPEc) colonies. Importantly, after exposure to gentamicin, the proportion of GAPEc colonies increased over time (***Figure 3B***), with petite colonies representing ∼ 90-100% of the total population at 24 h post-gentamicin exposure. Indeed, in PAP assays (***Figure 1F***), all colonies on 8 mg.L^−1^ gentamicin LB agar plates were the GAPEc morphotype. Furthermore, exposure to tobramycin and amikacin also resulted in petite *E. cloacae* colonies whose proportion increased over time (***Figure 3B***). Importantly, neither aminoglycoside could completely sterilise *E. cloacae* cultures as the number of viable bacteria did not change between 5 and 24 h post-treatment (***Figure 3C***). These results indicate persistence-like behaviour of *E. cloacae* against clinically important aminoglycosides.

**Figure 3.**
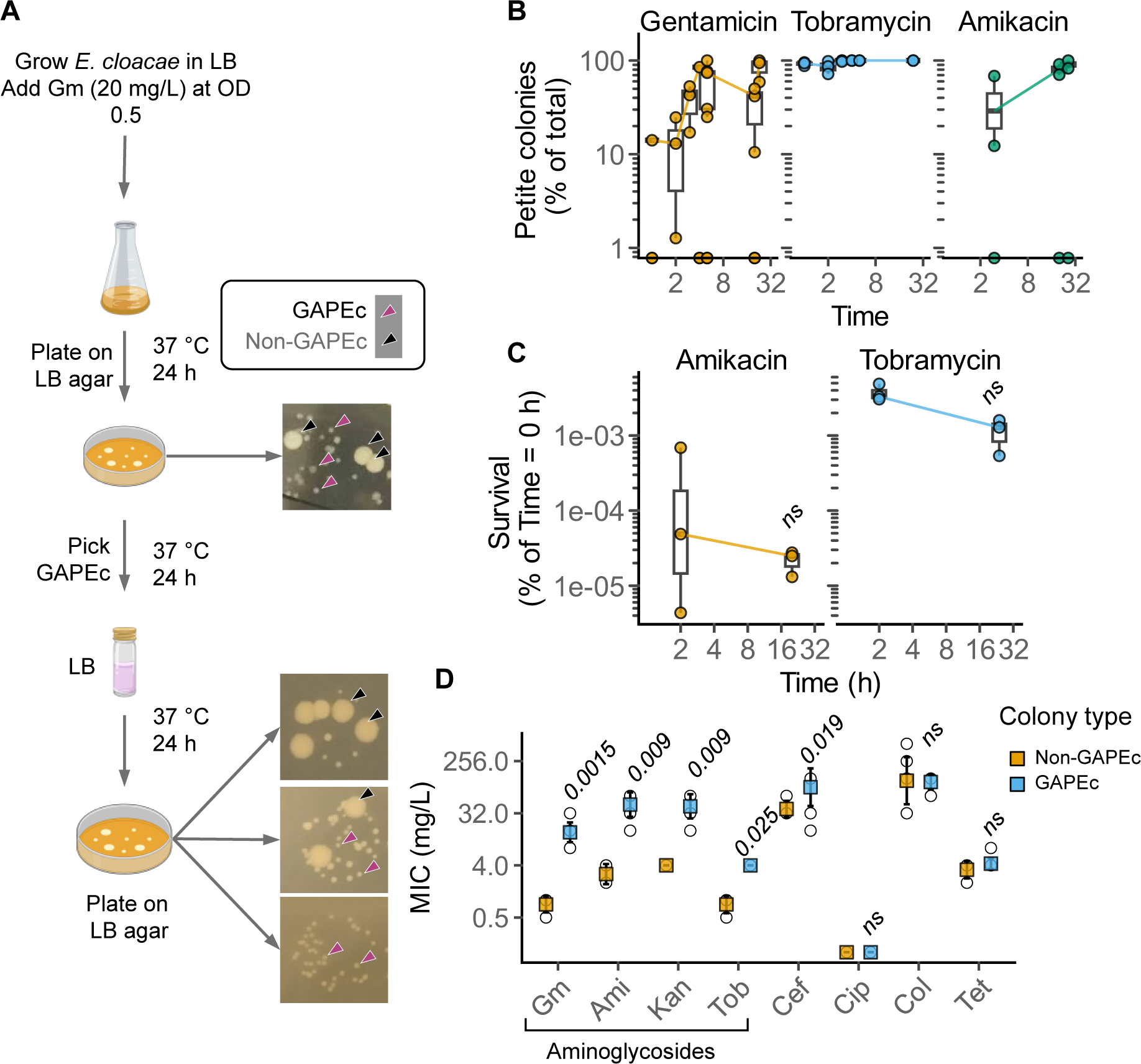
Aminoglycoside exposure of *E. cloacae* ATCC 13047 triggers petite colonies with increased MICs. **(A)** Schematic depiction of the appearance of gentamicin-associate petite *E. cloacae* (GAPEc) colonies after exposure of *E. cloacae* ATCC 13047 to gentamicin (20 mg.L^−1^), and their reversion back to ‘typical’ (non-GAPEc) colony morphotype after growth in broth without gentamicin. Arrows used to point to colony morphotypes indicated in the legend. **(B)** Percentage of petite colonies after exposure to the indicated aminoglycosides over time. *E. cloacae* cultures in exponential phase were treated with gentamicin (20 mg.L^−1^), tobramycin (20 mg.L^−1^) or amikacin (80 mg.L^−1^) for the times as indicated. Box and whisker with median shown from n = 3 independent experiments. **(C)** Survival of *E. cloacae* ATCC 13047 after exposure to amikacin (80 mg.L^−1^) or tobramycin (20 mg.L^−1^) at 5 or 24 h as indicated. Box and whisker with median shown from n = 3 independent experiments. ns = not significant for comparison of CFU at the two time points by mixed effects ANOVA. **(D)** MICs measured by broth microdilution for non-GAPEc and GAPEc colonies against gentamicin (Gm), amikacin (Ami), kanamycin (Kan), tobramycin (Tob), ceftriaxone (Cef), ciprofloxacin (Cip), colistin (Col) and tetracycline (Tet). Mean (coloured square) and SD shown from n = 8 (non-GAPEc) or n = 10 (GAPEc) independent experiments; open circles represent all data points. FDR-adjusted two-tailed *P* values for comparisons between GAPEc and non-GAPEc for each antibiotic from Mann-Whitney tests; ns = not significant (*P* > 0.05).

Importantly, however, the GAPEc phenotype was transient; petite colonies grown overnight in the absence of antibiotic produced both petite and normal sized colonies (***Figure 3A***). Taken together, we reason that GAPEc formation could enable bacteria to withstand aminoglycosides, and this subpopulation becomes enriched over time and forms a larger proportion of surviving bacteria.

### GAPEc have higher MICs for aminoglycosides

We hypothesised that heteroresistance arises because the GAPEc colony morphotypes have higher MIC against aminoglycosides. Indeed, MICs against gentamicin, amikacin, kanamycin and tobramycin, of GAPEc colonies were higher than non-GAPEc bacteria: ∼ 16-fold higher for gentamicin, ∼20-fold higher for amikacin, ∼12-fold higher for kanamycin, and ∼4-fold higher for tobramycin (***Figure 3D***). These results indicate that GAPEc have broad resistance to aminoglycosides with MICs higher than EUCAST breakpoints (2 mg.L^−1^). Interestingly, the MIC against ceftriaxone, a third-generation cephalosporin, also increased ∼4-fold (***Figure 3D***). In contrast, MICs of GAPEc against other antibiotic classes, including a fluoroquinolone (ciprofloxacin), a polymyxin (colistin) and tetracycline remained unchanged (***Figure 3D***). Overall, we conclude that the GAPEc morphotype promotes resistance mainly against aminoglycosides.

We next asked whether GAPEc morphotypes survive exposure to gentamicin for longer periods of time as compared to non-GAPEc ‘parental’ bacteria. Indeed, after exposure to 8 mg.L^−1^ gentamicin there was no change in the viability of GAPEc up to 2 h, whereas only ∼10^−3^ % non-GAPEc survived (***Figure 4A***). Similarly, ∼100-fold more GAPEc survived at 5 h than non-GAPEc (***Figure 4A***). We therefore conclude that GAPEc can withstand gentamicin without loss in viability for longer periods of time.

**Figure 4.**
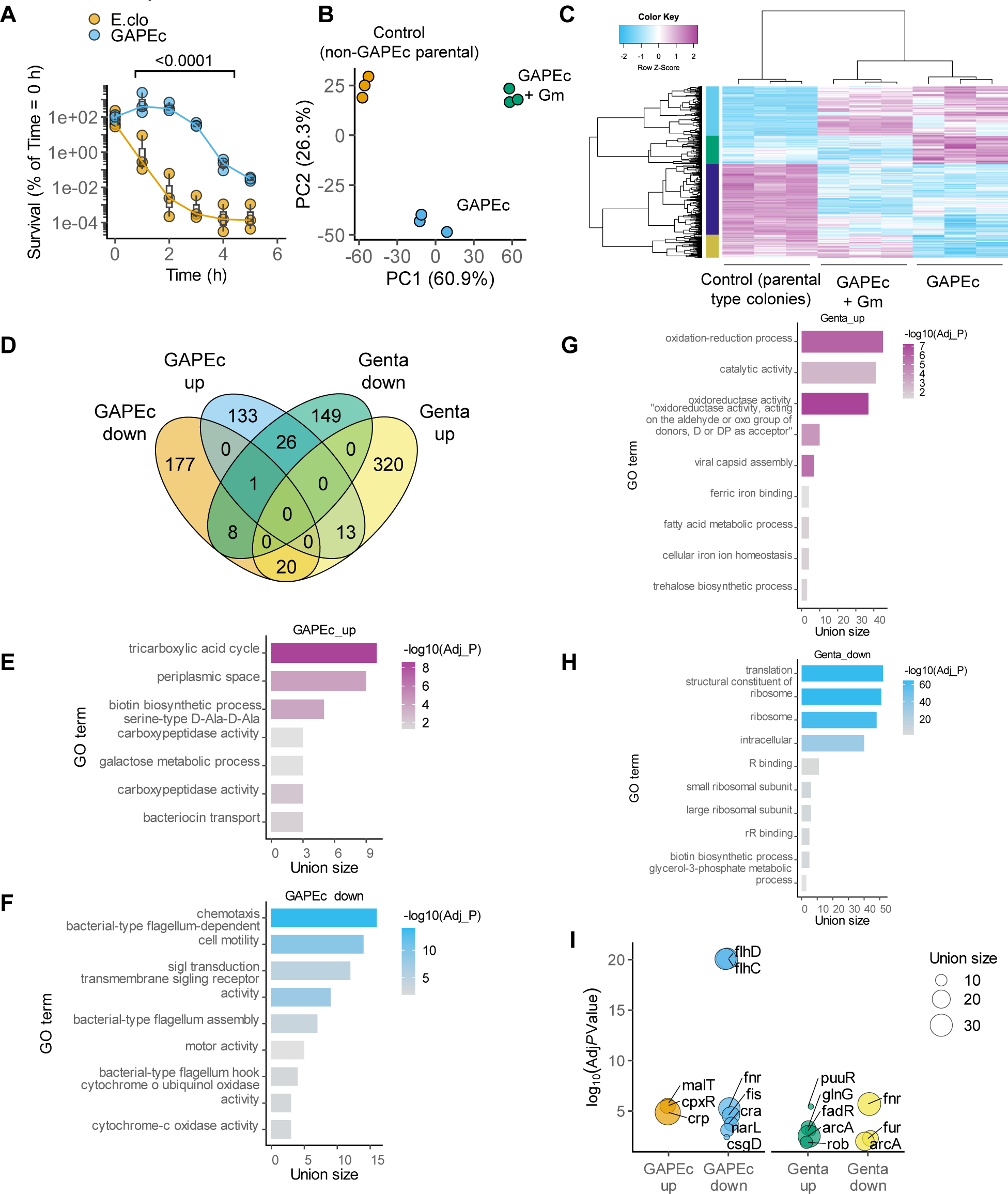
Transcriptional reprogramming drives heteroresistance to gentamicin. **(A)** Gentamicin time-to-kill curves after exposure to gentamicin (8 mg.L^−1^) for the indicated colony morphotypes of *E. cloacae* ATCC 13047. Box and whiskers with median plotted from n = 3 independent experiments. **(B-C)** Principal component analyses (**B**) and hierarchical clustering (**C**) of RNA sequencing data from the indicated growth conditions. Each dot in (B) represents an independent biological sample; n = 3 experiments. Colour key in (C) indicates raw Z-scores of normalised log2 counts per million for differentially expressed genes in the indicated conditions. Gm, gentamicin (8 mg.L^−1^). **(D)** Venn diagram showing the distribution of up- and down-regulated genes in the the indicated conditions following RNAseq analyses. **(E-H)** Gene ontology (GO) terms enriched in genes up-or down-regulated in the indicated comparisons. GAPEc = GAPEc vs non-GAPEc; Genta = GAPEc + gentamicin (8 mg.L^−1^, 2 h) vs GAPEc without gentamicin. The union size is the number of target genes in the data set. FDR-adjusted *P* values for GO terms plotted on -log_10_ scale. **(I)** Plot showing the statistically enriched transcription factors among differentially expressed genes as indicated. The union size is the number of target genes in the data set. Hits with FDR-adjusted *P* < 0.05 (Adj *P* Value) are plotted on a -log_10_ scale.

### Global gene expression changes in E. cloacae exposed to gentamicin

We next sought to determine the mechanisms that promote gentamicin resistance in GAPEc. Given the transient nature of the heteroresistance behaviour, we hypothesised that differential gene expression empowers GAPEc to resist gentamicin. To test this, we performed sequencing on mRNA from exponentially growing non-GAPEc (i.e., parental *E. cloacae* unexposed to gentamicin) and GAPEc (from 8 mg.L^−1^ LB-gentamicin plates in PAP assays). We also prepared RNA from exponentially growing GAPEc treated with gentamicin (8 mg.L^−1^) for 2 h, a time point at which there is no change in viable bacteria (***Figure 4A***). Consistent with our hypothesis, principal component analyses of RNA sequencing data revealed clear separation of the three treatment groups, indicative of distinct gene expression programmes in these conditions (***Figure 4B***). Unsupervised hierarchical clustering similarly resulted in neighbouring grouping of the three categories with many differentially regulated genes (***Figure 4C***). Overall, at log_2_(fold-change) ≥ 2 or ≤ −2 cut-offs, there were 377 differentially expressed genes when comparing GAPEc vs non-GAPEc, and 536 differentially expressed genes after treatment of GAPEc with gentamicin (***Figure 4D***; ***Supplementary Table S1-S2***). Notably, the highest upregulated genes in GAPEc were periplasmic proteases, chaperones, inner and outer membrane proteins, and maltose or trehalose transport genes; among the most downregulated were the flagellar apparatus and fimbrial biosynthesis genes (***Supplementary Table S1-S2***).

To identify metabolic processes and signalling pathways that could be responsible for gentamicin heteroresistance, we performed enrichment analyses on up- and downregulated genes. Among the upregulated genes in GAPEc were gene ontology (GO) terms representing periplasmic processes, the TCA cycle intermediates for supplying basic metabolic building blocks, biotin synthesis, galactose metabolism, and bacteriocin transport (***Figure 4E, Supplementary Table S3***). In contrast, processes such as chemotaxis and motility, and cytochrome c oxidase activity were downregulated in GAPEc (***Figure 4F, Supplementary Table S3***). Upon treatment with gentamicin, pathways involved in redox reactions and withstanding reactive oxygen species (ROS), iron uptake, and trehalose synthesis were enriched (***Figure 4G, Supplementary Table S3***). Not surprisingly, processes involving protein translation and ribosomal structure were markedly downregulated after gentamicin treatment (***Figure 4H, Supplementary Table S3***). Together, these analyses indicate that *E. cloacae* undergoes a broad shift in bioenergetics and substrate utilisation with a switch to anaerobic respiration, reduced motility, and altered processes in the periplasm and cell wall.

We next investigated transcription factors that might drive these gene expression changes. Because few datasets are available from direct studies on *E. cloacae,* we relied on cross-species gene mapping from studies in *E. coli* MG1655 (available on BioCyc) alongside manual curation. Notably, genes with the highest fold-change were targets of *cpxR* (ECL_05064), which is activated by its cognate sensor kinase *cpxA* (ECL_05065) in response to cell envelope stress (25) (***Figure 4I***). Targets of the global regulator such as *crp* (ECL_04734) and the maltose operon regulator *malt* (ECL_04784) were also upregulated, which indicated catabolite repression-like responses (***Figure 4I***). Among downregulated genes were many targets of *flhD* (ECL_01400) and *flhC* (ECL_01401), which is consistent with the reduced expression of the flagellar apparatus and chemotaxis genes (***Supplementary Table S1***). Additionally, *fnr* (ECL_02259), *csgD* (ECL_02600), *narL* (ECL_01619), *cra* (ECL_00876) and *fis* (ECL_04646) targets were enriched among downregulated genes in GAPEc. Gentamicin treatment led to enrichment of *puuR* (ECL_02228), *glnG* (ECL_05113), *fadR* (ECL_01510), *rob* (ECL_00809) and *arcA* (ECL_00811) targets among upregulated genes, whereas *fnr* (ECL_02259), *fur* (ECL_03032) and *arcA* (ECL_00811) targets were downregulated (***Figure 4I***). Taken together, we conclude that gentamicin exposure results in a cell broad activation of stress responses that promote anaerobiosis, reduced motility and adherence, and reduced cell division and growth.

### Cell envelope stress response is required for aminoglycoside heteroresistance

Several targets of the CpxRA stress response were among the most differentially regulated genes in GAPEc and upon further exposure to gentamicin (***Table 2***). Many CpxRA targets were differentially expressed in our RNAseq, including the upregulation of periplasmic chaperones (*spy*, *ppiA* and *cpxP*), proteases (*degP* and *htpX*), transporters (*acrD*, *btsT*, *yccA*), and downregulation of motility and chemotaxis proteins (*motA* and *tsr*) and iron acquisition (*efeU*) (***Table 2***).

**Table 2.** Differentially regulated genes predicted to be CpxRA targets.

S = sensitive; R = resistant, ND = not determined
| Condition | Gene ID | Common name | Product description | log2(FC) | Adj P value |
| --- | --- | --- | --- | --- | --- |
| GAPEc vs non-GAPEc | ECL_01450 | yebE | FIG004088: inner membrane protein YebE | 8.5 | 1.3E-07 |
|  | ECL_02444 | spy | Spheroplast protein Y | 6.4 | 5.8E-06 |
|  | ECL_05063 | cpxP | Periplasmic protein CpxP | 5.1 | 2.3E-07 |
|  | ECL_00965 | degP | HtrA protease/chaperone protein | 4.4 | 1.5E-07 |
|  | ECL_04004 | yqaE | UPF0057 membrane protein YqaE | 3.7 | 5.4E-05 |
|  | ECL_01472 | htpX | Protease HtpX | 3.3 | 9.2E-07 |
|  | ECL_03767 | acrD | Aminoglycosides efflux system AcrAD-TolC, inner-membrane proton/drug antiporter AcrD (RND type) | 3.2 | 3.2E-06 |
|  | ECL_04739 | ppiA | Peptidyl-prolyl cis-trans isomerase PpiA precursor (EC 5.2.1.8) | 2.9 | 1.2E-06 |
|  | ECL_00747 | btsT | Pyruvate:H<sup>+</sup> symporter BtsT | 2.7 | 1.5E-03 |
|  | ECL_02676 | yccA | Putative TEGT family carrier/transport protein | 2.7 | 2.6E-06 |
|  | ECL_04647 | yhdU | Uncharacterized protein YhdU | 2.6 | 1.7E-03 |
|  | ECL_00805 | slt | Soluble lytic murein transglycosylase (EC 4.2.2.n1) | 2.6 | 2.8E-05 |
|  | ECL_00561 | psd | Phosphatidylserine decarboxylase (EC 4.1.1.65) | 2.3 | 1.4E-04 |
|  | ECL_04140 | amiC | N-acetylmuramoyl-L-alanine amidase (EC 3.5.1.28) | 2.3 | 3.2E-05 |
|  | ECL_05122 | srkA | YihE protein, a ser/thr kinase implicated in LPS synthesis and Cpx signalling | 2.1 | 1.8E-04 |
|  | ECL_01131 | ampH | D-alanyl-D-alanine-carboxypeptidase/endopeptidase AmpH | 2.1 | 7.1E-06 |
|  | ECL_00917 | yacH | Uncharacterized protein YacH | 2.0 | 1.6E-05 |
|  | ECL_00152 | gpmM | 2,3-bisphosphoglycerate-independent phosphoglycerate mutase (EC 5.4.2.12) | -2.6 | 6.1E-05 |
|  | ECL_01867 | yncD | Probable tonB-dependent receptor yncD precursor | -2.7 | 1.7E-03 |
|  | ECL_02615 | ECL_02615 | Ferrous iron transport permease EfeU | -2.8 | 6.9E-04 |
|  | ECL_00760 | tsr | Methyl-accepting chemotaxis protein I (serine chemoreceptor protein) | -2.9 | 8.3E-07 |
|  | ECL_01189 | cyoA | Cytochrome O ubiquinol oxidase subunit II (EC 1.10.3.-) | -2.9 | 1.1E-03 |
|  | ECL_01402 | motA | Flagellar motor rotation protein MotA | -4.5 | 1.1E-06 |
| Gentamicin vs w/o Gentamicin | ECL_01053 | fadE | Acyl-coenzyme A dehydrogenase FadE (EC 1.3.8.-) | 5.3 | 7.4E-06 |
|  | ECL_02449 | astC | Succinylornithine transaminase (EC 2.6.1.81) | 5.1 | 6.2E-05 |
|  | ECL_00330 | acs | Acetyl-CoA synthetase (EC 6.2.1.1) | 4.4 | 6.3E-05 |
|  | ECL_01604 | chaB | Cation transport regulator chaB | 4.0 | 3.2E-04 |
|  | ECL_01392 | ftnB | Bacterial non-heme ferritin (EC 1.16.3.2) | 2.7 | 2.3E-04 |
|  | ECL_01394 | araF | L-arabinose ABC transporter, substrate-binding protein AraF | 2.4 | 3.8E-04 |
|  | ECL_03683 | fadI | 3-ketoacyl-CoA thiolase (EC 2.3.1.16) | 2.2 | 8.1E-05 |
|  | ECL_04646 | fis | DNA-binding protein Fis | -2.2 | 2.2E-04 |
|  | ECL_02999 | cydA | Cytochrome d ubiquinol oxidase subunit I (EC 1.10.3.-) | -2.2 | 3.6E-04 |
|  | ECL_04129 | sdaC | Serine transporter | -2.6 | 1.2E-03 |
|  | ECL_01189 | cyoA | Cytochrome O ubiquinol oxidase subunit II (EC 1.10.3.-) | -2.7 | 1.8E-03 |
|  | ECL_04329 | exbB | TonB-ExbBD energy transducing system, ExbB subunit | -4.0 | 1.1E-05 |
|  | ECL_00747 | btsT | Pyruvate:H<sup>+</sup> symporter BtsT | -5.6 | 1.8E-05 |

The response regulator CpxR (ECL_05064) is activated upon phosphorylation by the CpxA (ECL_05065) sensor histidine kinase that is activated by altered periplasmic homeostasis and phosphorylation (25). Interestingly, CpxA may phosphorylate other response regulators, and on the other hand, CpxR can be activated by other sensor kinases (25, 26). Therefore, to investigate the role of the CpxRA two-component system, we generated a scarless Δ*cpxRA E. cloacae* strain lacking both genes. Remarkably, PAP assays showed that the Δ*cpxRA* strain completely lost heteroresistance to gentamicin (***Figure 5A***), pointing to an indispensable role for cell envelope stress response in this process. We complemented the Δ*cpxRA* strain by introducing a single copy of *cpxRA* under their natural promoter (Δ*cpxRA*+CpxRA); a strain containing a single copy of mVenus fluorescence protein under the ‘Ultra’ promoter (27) served as a negative control (Δ*cpxRA*+Ven). Lack of heteroresistance in Δ*cpxRA* bacteria was restored to wild-type levels in the Δ*cpxRA*+CpxRA strain (***Figure 5A***); the Δ*cpxRA*+Ven strain behaved similarly to the scarless Δ*cpxRA* strain indicating that insertion of genes at the attTn7 site downstream of the *glmS* locus did not have adverse effects in *E. cloacae* (***Figure 5A***). All strains grew similarly in LB, which ruled out viability defects under standard growth conditions (***Figure 5B***). Together, these results conclusively show that the cell envelope stress response driven by the CpxRA regulon is indispensable for gentamicin heteroresistance in *E. cloacae*.

**Figure 5.**
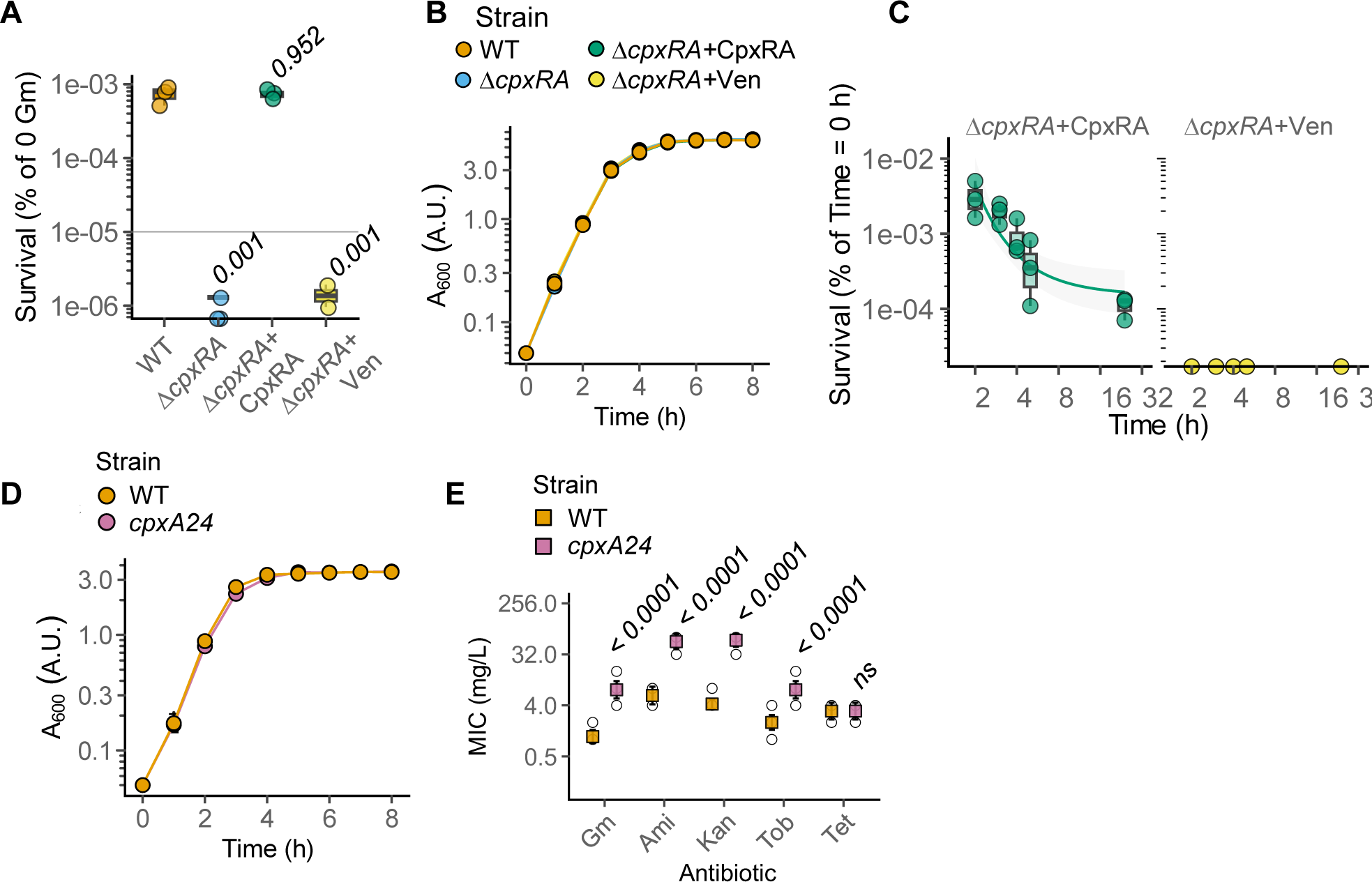
The CpxRA cell envelope stress response pathway is required for gentamicin heteroresistance. **(A)** PAP assay showing proportion of surviving bacteria of the indicated *E. cloacae* ATCC 10347 genotypes on agar plates with 8 mg.L^−1^ gentamicin. Box and whisker with median shown from n = 3 independent experiments. Two-tailed *P* values for the comparisons with wild type (WT) *E. cloacae* from mixed effects ANOVA. **(B)** Growth curves of the indicated strains *E. cloacae* ATCC 13047 strains plotted as mean ± SD from n = 3 experiments. **(C)** Gentamicin time-to-kill curves for the indicated complemented and Δ*cpxRA* knockout *E. cloacae* ATCC 13047 showing the percentage survival over time after exposure to gentamicin (8 mg.L^−1^). Graph shows first order decay curve fit to data. Box and whiskers with median are plotted from n = 3 independent experiments. **(D)** Growth curves of the indicated *E. cloacae* ATCC 13047 WT and *cpxA24* plotted as mean ± SD from n = 3 experiments. **(E)** MICs measured by broth microdilution for *E. cloacae* ATCC13047 WT and *cpxA24* strains against gentamicin (Gm), amikacin (Ami), kanamycin (Kan), tobramycin (Tob) and tetracycline (Tet). Mean (coloured square) and SD shown from n = 12 independent experiments. Open circles represent all data points. FDR-adjusted two-tailed *P* values of comparisons between WT and *cpxA24* are shown from non-parametric ANOVA following aligned-rank transformation.

We next asked whether the loss of *cpxRA* also sensitised *E. cloacae* to rapid and complete killing by gentamicin. Indeed, the Δ*cpxRA+*Ven strain was rapidly killed within 1 h with no detectable CFU after 1 h post-treatment (***Figure 5C***); the Δ*cpxRA+*CpxRA behaved similarly to the wild-type strain displaying a biphasic kill-curve (***Figure 5C***) similar to that seen with the wild-type strain (***Figure 1C***). Taken together, we conclude that the CpxRA two-component system is essential for phenotypic resistance to gentamicin in *E. cloacae*.

We next asked whether constitutively active envelope stress response can recapitulate these phenotypes. In *E. coli* the *cpxA24* allele lacking 32 aa in the periplasmic loop of the sensor kinase CpxA is constitutively active (25). As *E. cloacae* CpxA (ECL_05065) is > 95 % identical in amino acid sequence to the *E. coli* CpxA (***Figure S1***), we generated a similar deletion and introduced this allele (*cpxA24*) at the native genomic locus in *E. cloacae*. Wildtype and *cpxA24 E. cloacae* grew similarly in normal growth conditions, ruling out broad defects in growth-rates in standard media (***Figure 5D***). Notably, *E. cloacae cpxA24* also had higher MICs for aminoglycosides, but not tetracycline (***Figure 5E***), which is similar to the GAPEc morphotype. Together with the results on GAPEc, these findings in *E. cloacae cpxA24* establish a pivotal role for the cell envelope stress response in aminoglycoside resistance.

### Copper- and CpxRA-driven aminoglycoside resistance in E. cloacae strains

We reasoned that as GAPEc bacteria have high expression of the envelope stress regulon, heterologous activation of this pathway would also result in higher heteroresistance. The CpxRA pathway is activated in response to envelope stress by environmental factors, including Cu^2+^, which is an antimicrobial heavy metal (25). Consistent with this, *E. cloacae* exposed to 4 mM Cu^2+^ had ∼6-16—fold higher gentamicin MIC (***Figure 6A***); whereas the MIC of the Δ*cpxRA* strain did not increase above the breakpoint (2 mg.L^−1^) (***Figure 6A***). Notably, both strains had a similar Cu^2+^ MIC of 16 mM, indicating that *cpxRA* are dispensable for copper-resistance in *E. cloacae* (***Figure 6B***). Taken together, we show that copper exposure confers aminoglycoside resistance in a *cpxRA*-dependent manner.

**Figure 6.**
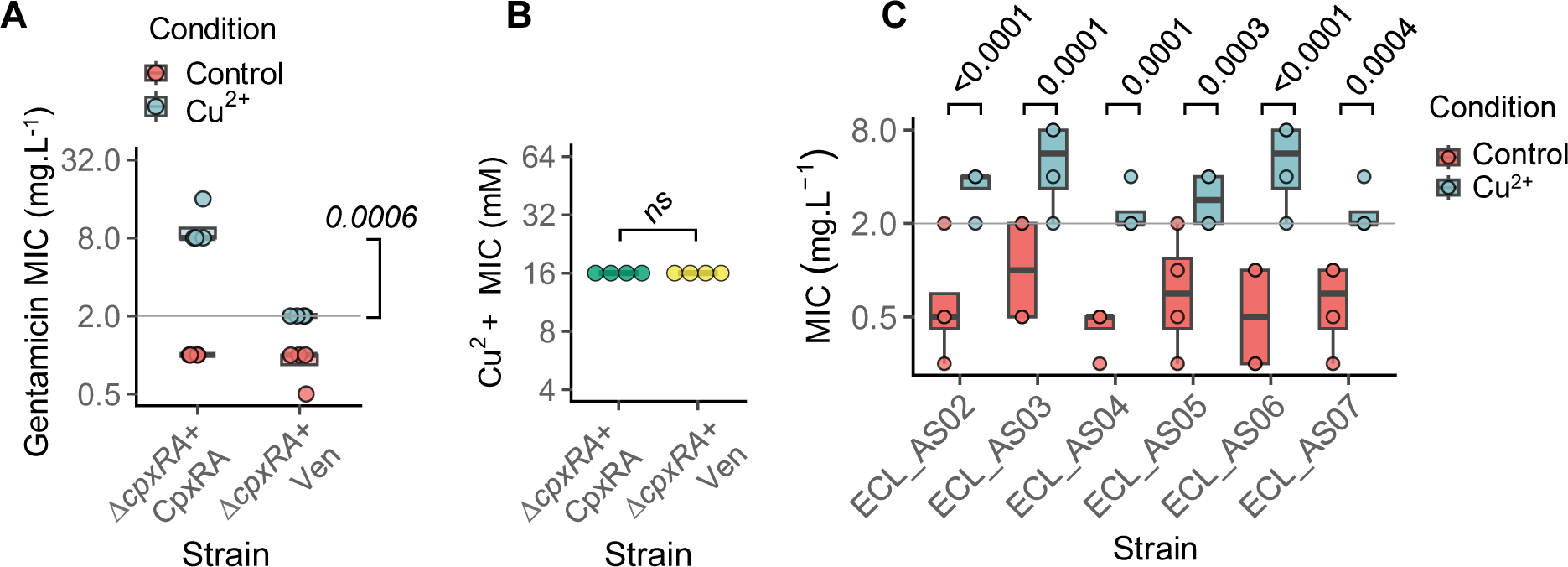
Exposure to copper increases gentamicin resistance via *cpxRA*. **(A)** Gentamicin MIC measured by broth microdilution for the indicated complemented or Δ*cpxRA* knockout *E. cloacae* ATCC 13047. MICs were measured in the absence or presence of 4 mM copper sulphate. Box and whisker with median shown from n = 4 independent experiments. FDR-adjusted two-tailed *P* value for comparison between the MIC of the two genotypes in the presence of Cu2+ from non-parametric ANOVA following aligned-rank transformation. **(B)** Cu^2+^ MIC measured by broth microdilution for the indicated complemented or Δ*cpxRA* knockout E. cloacae ATCC 13047. Box and whisker with median shown from n = 4 independent experiments. ns, not significant, i.e., two-tailed *P* value > 0.05 for the indicated comparisons from Mann-Whitney test. **(C)** Gentamicin MICs for the indicated clinical E. cloacae isolates in the absence (“Control”) or presence of Cu^2+^ (4 mM copper sulphate). Box and whiskers blot with median from n = 4 independent experiments. Grey line indicates EUCAST break point. FDR-adjusted two-tailed *P* values are shown for indicated comparisons in the presence of Cu^2+^ for each strain from non-parametric ANOVA following aligned-rank transformation.

We further hypothesised that exogenously increasing *cpxRA* with copper would lead to increased gentamicin MIC in the clinical isolates. We measured the gentamicin MICs of all 6 clinical strains in the absence or presence of 4 mM Cu^2+^. Notably, similar to the ATCC 13047 type-strain, copper increased gentamicin MICs for all 6 clinical isolates to levels above the EUCAST breakpoint (***Figure 6C***). We therefore conclude that copper- and *cpxRA*-dependent enhancement of aminoglycoside resistance is not just a trait of the laboratory strain but also seen in sepsis-related isolates from patients. Altogether, our results establish that *E. cloacae* strains display phenotypic aminoglycoside resistance that can be enhanced by envelope stress, for example through exposure to copper.

## Discussion

Here we established that *E. cloacae* contains an aminoglycoside resistant subpopulation that arises through the activation of the CpxRA cell envelope stress response and correlates with a petite colony morphotype. In all *E. cloacae* strains tested we tested, copper, an inducer of the CpxRA stress regulon, increased aminoglycoside resistance, pointing to its broad conservation in this important group of environmental and hospital-associated bacteria (28, 29). CpxRA has been implicated in fosfomycin resistance in *E. coli* (30, 31) and colistin resistance in *Salmonella enterica* serovar Typhimurium (32). Mutations in *cpxRA* have been linked to antibiotic resistance in clinical isolates of important Gram-negative bacteria, including *K. pneumoniae* (33), *Klebsiella aerogenes, Salmonella enterica* serovar Typhi (34), and *Proteus mirabilis* (35). In *Acinetobacter baumanii,* aminoglycoside heteroresistance results via the amplification of aminoglycoside resistance genes (36). However, here we report that differential gene expression via CpxRA can also confer heteroresistance in sensitive bacteria. These findings add to the interest in developing CpxA kinase inhibitors for combination therapy along with conventional antibiotics (37).

We observed both persistence- and heteroresistance-like behaviour across different strains of *E. cloacae*; importantly, complete loss of viable bacteria was not observed in any experiment at 24 h post-treatment. Experimental variability is not surprising in such studies and have been previously reported (8, 38, 39); it is plausible that this is due to the small size of the resister subpopulation which could result in low detection rates or require longer incubation time to detect growth of the surviving subpopulation. Importantly, serum concentrations of gentamicin peak at 20 mg.L^−1^ and reduce to < 8 mg.L^−1^ after ∼ 5 h post-administration (40). Our results indicate that these concentrations would not only fail to sterilise the *E. cloacae* type-strain or clinical isolates (***Figures 1D, 2B***) but may also facilitate heretoresistant growth. Likewise, peak amikacin and tobramycin plasma concentrations range from 10-60 mg.L^−1^ and <10 mg.L^−1^, respectively (41–43), neither of which is sufficient to sterilise *E. cloacae in vitro*. The increase in ceftraixone MIC in GAPEc is explained by the upregulation of the β-lactamase *ampC* (***Figure 3D***, ***Supplementary Table S1-S2***), presumably due to cell envelope stress. Increased cross-resistance to ceftriaxone is also a concerning observation given the use of cephalosporins and aminoglycosides in combination therapy.

The sensor kinase CpxA phosphorylates the response regulator CpxR, which was originally identified as a conjugative pilus expression regulator (25). CpxR regulates ∼90 genes in *E. coli*, many of which include proteases (e.g., DegP), chaperones (e.g., DsbB, PpiA), redox enzymes, transporters, and cell wall-modifying enzymes, which together rectify envelope stress (25). CpxRA can suppress chemotaxis and motility, increase adherence and drug efflux, which our RNA sequencing show is also conserved in *E. cloacae*. Transcriptional reprogramming in GAPEc leads to a shift in metabolism to anaerobic respiration and increases catabolic activity to reduce the proton-motive force, which is required for aminoglycoside uptake (16). Anaerobiosis was also evident from reduced expression of electron transport chain and upregulation of predicted targets of fumarate nitrate reductase regulator (FNR) and nitrate/nitrite response regulator (NarL) (44), whereas starvation- and catabolite repression-like responses are suggested from the signature of cyclic-AMP response protein (Crp)-regulated genes (45). The upregulation of key efflux pumps (e.g., AcrAD) could facilitate gentamicin efflux. Enrichment of various membrane proteins and the upregulation of maltose and trehalose transporters is suggestive of an imbalance in osmolytes in the face of envelope stress. Enzymes that counteract damage from ROS were also upregulated upon gentamicin exposure. We conclude that the CpxRA pathway drives a protective response that reduces growth and aerobic respiration and promotes protein re-folding and/or degradation to protect against aminoglycosides. Future work should focus on why heteroresistance was only observed in exponentially growing *E. cloacae* and whether CpxRA activation is more robust in actively growing bacteria. The precise cause of cell envelope stress that triggers CpxA after exposure to aminoglycosides also remains known. It is plausible that aminoglycosides bind and disrupt the LPS outer membrane. Alternatively, aminoglycoside-induced error-prone translation could result in the accumulation of mistranslated or unfolded proteins leading to a breakdown in periplasmic homeostasis.

Typically, ‘small colony variants’ (SCVs) are observed in response to auxotrophy (46), e.g., to menadione, thiamine or heme, or mutations in the electron transport chain in *Staphylococcus aureus* (47). The slow growth rate of SCVs confers stable resistance to many classes of antibiotic, including aminoglycosides, trimethoprim-sulphamethoxazole, tetracycline, ciprofloxacin, oxacillin, clindamycin and daptomycin (47). SCVs are also known in Gram-negative bacteria, e.g., *Salmonella enterica, E. coli, Shigella, Pseudomonas, Klebsiella, Neisseria, Brucella, Serratia, E. cloacae* (46, 48–50). In contrast, here we describe transient petite colony and high aminoglycoside resistance phenotypes that readily revert to ‘typical’ *E. cloacae* behaviours. To the best of our knowledge, this is the first report linking CpxRA stress response to petite colony morphotype.

Copper is well-known for its antimicrobial effects and triggers the CpxRA stress response through ROS-induced protein misfolding or mislocalisation of outer membrane porins (25, 51). There is much interest in developing copper-coated surfaces and nanoparticles, and copper-based antimicrobials (51, 52). Therefore, our observation that copper increased gentamicin resistance to > 8 mg.L^−1^ in *E. cloacae* is worrying, especially because gentamicin treatment regimens can be ineffective against bacteria with an MIC > 2 mg.L^−1^ (53). Many opportunistic pathogens, including *E. cloacae*, often carry plasmids that encode genes for heavy-metal resistance, for instance through co-selection in the presence of heavy metals and antibiotics in effluents in their environment (54). Multiple homologues of the copper sensing histidine kinase CusS are present in the *E. cloacae* genome. The ATCC 13047 strain additionally carries copper-resistance genes on the pECL_A plasmid (21); this likely explains the similar sensitivity to copper of the wild-type and Δ*cpxRA E. cloacae*.

A strength of our work is the use of diverse clinical bloodstream isolates from an ongoing cohort study of patients admitted to hospital from the community with suspected sepsis (22). These isolates are from epidemiologically unrelated patients and are not biased by specific characteristics often required for archiving, such as antimicrobial susceptibility or any association with outbreaks (22). Gentamicin, amikacin and tobramycin are the most commonly used aminoglycosides in the clinic, including against multidrug resistant Gram-negative infections and are more efficacious and better tolerated than colistin and tigecycline (55). Aminoglycosides are generally administered intravenously in hospital settings, although topical or oral neomycin is also used against skin infection (56, 57). Aminoglycosides do have downsides, such as nephrotoxicity and ototoxicity, however these can be managed through appropriate dosing (56, 57). The newest aminoglycoside, plazomicin, was approved in 2018, pointing towards the continued interest in this class of broad-spectrum bactericidal antibiotics (58). Therefore, guarding against phenotypic aminoglycoside resistance important is critical. As the CpxRA operon can be triggered by heavy metals (e.g., copper, iron), our results also suggest caution against the use of heavy metals as antimicrobial surface coatings to avoid inadvertent aminoglycoside heteroresistance.

## Acknowledgements

This manuscript is dedicated to the memory of Nicholas Lim. AJC, HS, HL, JB, HWY, NL were students on the MRes in Bacterial Pathogenesis and Infection and acknowledge support from the programme team. The authors thank Ewurabena A. Mills for help. *E. coli* MG 1655 was a gift from Alex McCarthy (Imperial College London), and MFDpir from Jean-Marc Ghigo (Institut Pasteur, Paris). The BioAID (Bioresource in Adult Infectious Disease) cohort study supported by the Imperial National Institute of Health Research Comprehensive Biomedical Research Centre (REC ref: 14/SC/0008).

## Author contributions

Investigation: AJC, DJB, EK, HS, HL, JB, HWY, NL, VM, VM, AC-P; Validation: JB, DJB, HS, HL, EK; Visualisation: AJC, DJB, EK, HL, HS, ARS; Writing – original draft: AJC, ARS; Writing – review and editing: AJC, DJB, JB, VM, AC-P, ARS; Supervision: VM, AC-P, DJB, ARS; Software: ARS; Resources: FD, SS; Formal analysis: AJC, DJB, HS, ARS; Conceptualisation: ARS; Funding acquisition: ARS, SS.

## Materials and Methods

### Ethics statement

The BioAID (Bioresource in Adult Infectious Disease) cohort study (UK CRN reference: 36653) was approved by a national research ethics committee REC ref: 14/SC/0008.

### Source of strains and antibiotics

All strains used in this study are listed in ***Tables 1* and *3***. Antibiotics are listed in ***Table 4***.

**Table 3.**
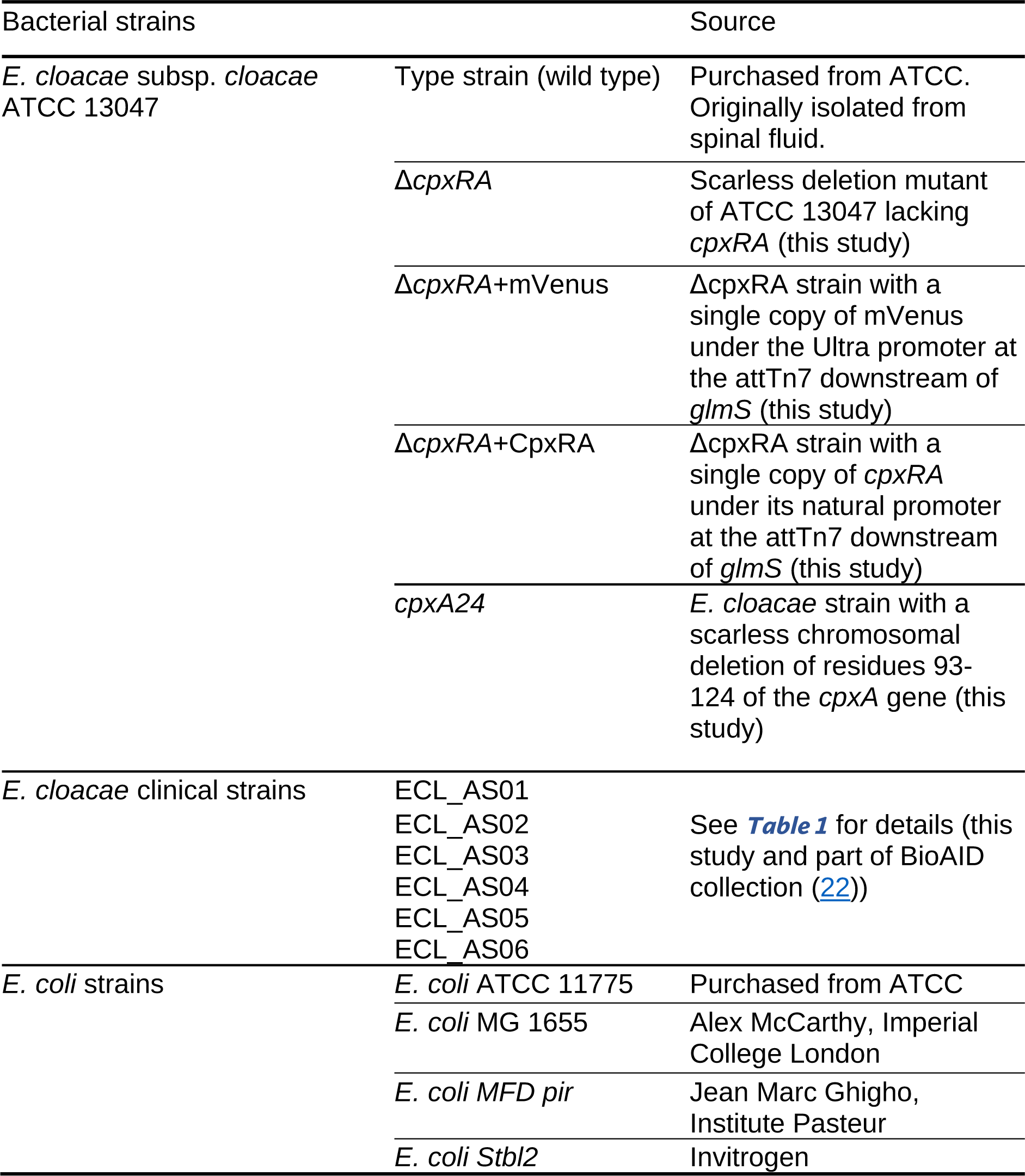
Bacterial strains used in this study.

**Table 4.** Sources of antibiotics used in this study.

| Product name | Product code, Company |
| --- | --- |
| Ampicillin Sodium Salt | 10419313, Fisher Scientific |
| Amikacin Sulphate Salt | 361958, Fluorochem |
| Ceftriaxone sodium hemiheptahydrate | C2226, Tokyo Chemical Industry |
| Ciprofloxacin | 17850, Sigma-Aldrich |
| Colistin Sulphate | C5647, LKT Laboratories |
| Gentamicin Solution | G1272, Sigma-Aldrich |
| Kanamycin Sulphate<br>from *Streptomyces kanamyceticus* | K4000-25G, Sigma-Aldrich |
| Streptomycin Sulphate Salt | S9137-25G, Sigma-Aldrich |
| Tetracycline | 87128, Sigma-Aldrich |
| Tobramycin | BS-9878A-1G, Molecular Dimensions |

### E. cloacae strains and preparation of starting cultures for all experiments

*E. cloacae* ATCC 13047 and *E. coli* ATCC 11775 were obtained from ATCC (***Table 2***), and clinical isolates were collected under the BioAID study (***Table 1***). Starter cultures (i.e., 5 mL overnight cultures) for experiments were prepared from frozen DMSO-stocks as recommended for studies on *E. coli* persisters to reduce variability across experiments and ensure inoculation of similar bacterial numbers into experimental culture flasks (38). Briefly, DMSO-stocks were prepared as follows: a single *E. cloacae* colony (type strains or clinical isolates) from a freshly streaked out LB agar plate was inoculated into 5 mL filter-sterilised lysogeny broth (LB; L3022, Sigma-Aldrich) with ampicillin (100 µg.mL^−1^) and incubated at 37 °C with shaking (180 rpm). This culture was diluted 1:100 into fresh 50 mL LB prepared in a 250 mL flask and grown at 37 °C with shaking to an optical density 600 nm (OD_600_) of 0.5, following which 9.2 mL culture was mixed with 0.8 mL DMSO, and 100 µL aliquots were prepared in sterile screw-capped tubes and stored at −80 °C. Overnight cultures for all experiments were prepared by inoculating 50 μL from DMSO stocks into 5 mL LB to have a uniform number of starting bacteria in the pre-inoculum. OD_600_ of 0.1 was found to correspond to ∼10^8^ CFU.mL^−1^ for *E. cloacae* by plating.

### Measurement of bacterial antibiotic susceptibility

Minimum inhibitory concentrations (MIC) were determined by the broth microdilution method ^42^. Mueller-Hinton Broth (MHB; 70192, Sigma-Aldrich, Spain) was autoclaved and cation-adjusted (with filter-sterilised 1000x stock of MgCl_2_ and CaCl_2_) to a final concentration of 10 mg.mL^−1^ Mg^2+^ and 20 mg.mL^−1^ Ca^2+^. Stationary phase cultures (37°C, 18-24 h, with shaking) were washed twice with sterile phosphate buffered saline (PBS; D8537, Sigma-Aldrich) and diluted to 5 x 10^5^ CFU.mL^−1^ for experiments. Two-fold dilutions of antibiotics were used (technical duplicates) in round-bottom 96-well plates, including wells without antibiotics and wells without bacteria as controls. Plates were incubated in a humidified 37 °C chamber for 24 h before noting the lowest antibiotic concentration that inhibited visible bacterial growth. MIC breakpoints from EUCAST were used to classify bacteria as susceptible or resistant to each antibiotic tested, whose sources are listed in ***Table S3***. These were 0.5 mg.L^−1^ for ciprofloxacin, 2 mg.L^−1^ for gentamicin, tobramycin, ceftriaxone, colistin; 4 mg.L^−1^ for tetracycline; 8 mg.L-1 for amikacin and kanamycin.

For the disk-diffusion method, 100 μL of 10^8^ CFU.mL^−1^ bacterial culture was spread on MHB agar plates and disks containing 10 µg gentamicin were placed on the agar surface within 15 min. Disks were prepared based on EUCAST guidelines. Plates were incubated overnight at 37 °C and the diameter of the zone of inhibition was measured and compared to the EUCAST zone diameter breakpoint (17 mm for gentamicin for *Enterobacterales*) (23).

### Aminoglycoside time-to-kill experiments

Mid-exponential phase (OD_600_ 0.5) cultures (prepared from overnight cultures started with frozen stocks as above) were treated with 20 mg.L^−1^ gentamicin, 80 mg.L^−1^ amikacin or 20 mg.L^−1^ tobramycin and samples collected at various time-points, washed twice in sterile PBS before preparing serial dilutions for enumerating viable bacterial counts. Typically, 3-, 5-or 10-fold serial dilutions were prepared, of which 10 μL were spotted onto agar plates (2 technical replicates), and spots with 3-50 colonies counted and a mean CFU.mL^−1^ calculated per experiment (detection limit: 10^2^ CFU.mL^−1^).

### Population Analysis Profile (PAP) assays

PAP assays were performed to detect heteroresistance as described before (24, 59). Bacteria were grown overnight from DMSO stocks, inoculated at 1:100 dilution in fresh 50 mL LB in a 250 mL flask and at OD = 0.5, 10 – 50 mL culture was centrifuged (5000 x*g*, 5 min), washed once in sterile PBS, and serial 3-, 5-or 10-fold dilutions as appropriate were prepared in PBS and duplicate 10 μL spotted on agar plates containing 0, 2, 4, 8 or 16 mg.L^−1^ gentamicin. Plates were incubated overnight at 37 °C and the proportion of surviving bacteria at each concentration of gentamicin relative to gentamicin-free plate was calculated. As the MIC for *E. cloacae* is ∼1 mg.L^−1^, heteroresistance was defined as the appearance of viable bacteria at a frequency of ≥ 10^−7^ on plates containing 8 mg.L^−1^ gentamicin. The CFU detection limit was 10^2^ CFU.mL^−1^, and experiments used ≥ 10^10^ bacteria to have sufficient sensitivity to detect heteroresistance. For bacterial cultures treated with gentamicin for 24 h, which contain very low CFU.mL^−1^, 100 mL culture was centrifuged, washed twice, and used for PAP assays.

### Molecular cloning and plasmids

During our studies, we found that *E. cloacae* carrying the J23100-mRFP-331Bb (Addgene plasmid #78271, gift from Tom Ellis (60)) which confers chloramphenicol resistance and drives mRFP expression using a strong, synthetic promoter J23100 and ribosome binding site (RBS) were poorly fluorescent. Similar lack of expression was observed with the pA1 promoter in another plasmid (not shown). The “Ultra” promoter from the pUltra series of plasmids (that confer kanamycin resistance (27)) was used to generate pJC-Ultra plasmid expressing mTurquoise2 (from pLifeAct-mTurquoise2; Addgene plasmid #36201, gift from Dorus Gadella (61)) in the 331Bb backbone, which led to detectable fluorescence. However, for reasons not yet clear, exposure to gentamicin and petite colony formation led to plasmid loss. We therefore developed a plasmid that integrates at the T7 insertion site 3’ of *glmS* through homologous recombination. The conjugative pTOX6 vector (62) (R6Kγ origin of replication) was modified for this purpose by removing the rhamnose regulator (digested with blunt-end cutters StuI and BstZI17I enzymes and re-ligated) and then the Tse2 toxin (digested with FspI & HincII enzymes and re-ligated) to generate pTraCam (which retains *tra* genes for conjugative transfer and Cam^R^). Cloning of the Ultra-promoter-RBS-mTrq2 into PstI cut pTraCam via sequence and ligase independent cloning (SLIC; (63)) generated the pTraCam-Ultra-mTrq2 plasmid. A region (∼500 bp) flanking the T7 insertion 3’ of *glmS* was cloned into the XhoI site of pTraCam-Ultra-mTrq2 to generate pTraCam-Ultra-mTrq2-EcloGlmS. All cloning steps were carried out in MFDpir *E. coli* (64), which was also used to conjugate this plasmid into *E. cloacae.* Conjugants were confirmed as chloramphenicol resistant and the presence of natural pECL_A and pECL_B plasmids was confirmed by PCR. Proof-reading polymerases such as Phusion or KOD were used for PCR cloned fragments were verified by Sanger sequencing (Azenta). All primers used in this study are listed in ***Table 5***.

**Table 5.**
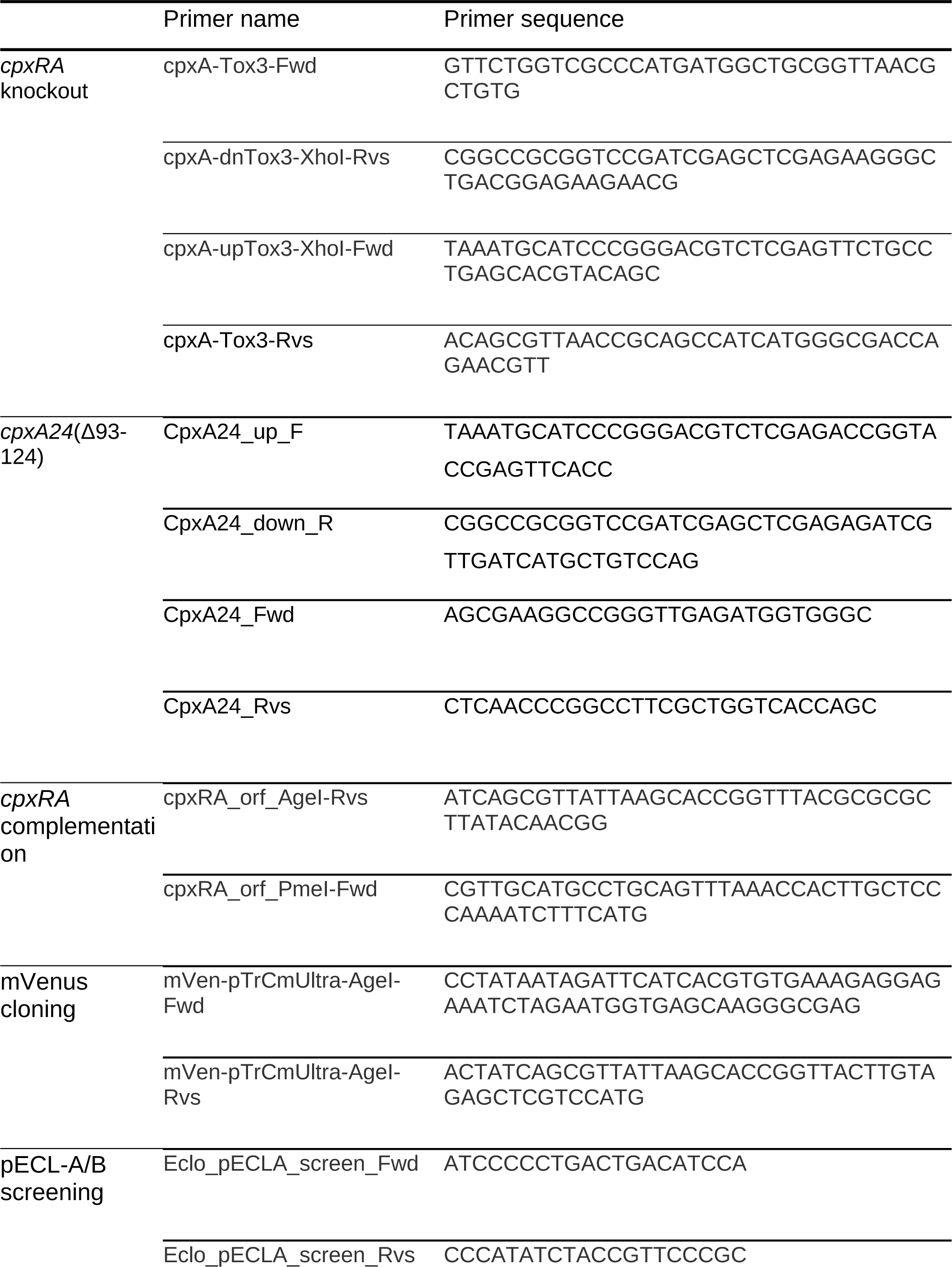
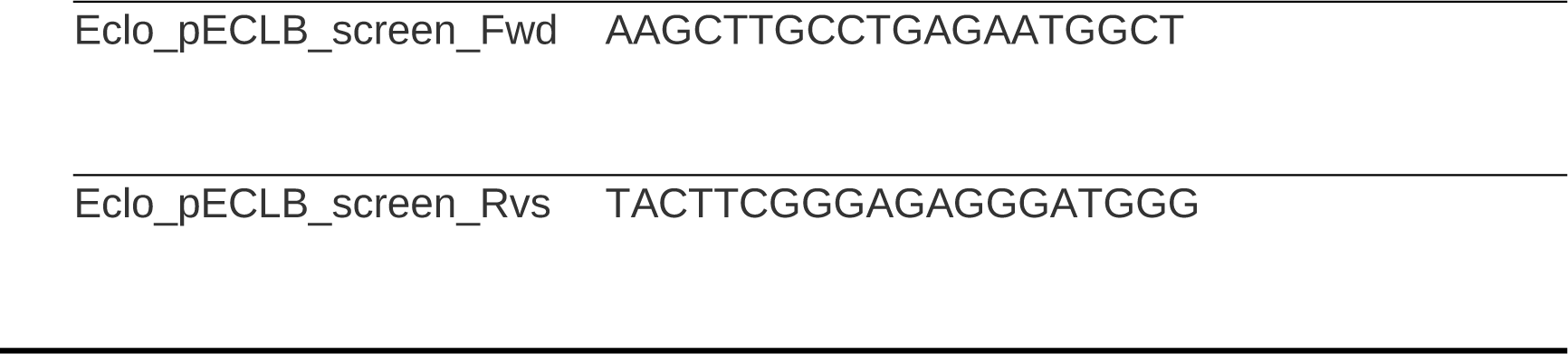
Primers used in this study.

### Generation of E. cloacae *cpxA* knockout, *cpxA24* and complemented strains

Scarless deletion of the *cpxRA* locus in *E. cloacae* ATCC 13047 used a two-step process relying on the pTOX6 plasmid system (62). Approximately 500 bp flanking regions were cloned to allow homologous recombination and deletion of most of the open reading frame of CpxRA. The KOD hot-start polymerase was used to generate flanking fragments which were cloned in a 3-way SLIC into pTOX6 using *E. coli* MFDpir as the host with all steps including glucose (2% w/v) and diaminopimelic acid (DAP; 0.3 mM). Single crossovers were obtained by conjugation into *E. cloacae* with 3:1 ratio of overnight cultures of donor *E. coli* and recipient *E. cloacae* plated in a 100 μL spot on LB agar for 4 h at 37°C, followed by collecting colonies, washing in 1 mL PBS and plating serial dilutions on LB agar containing glucose (2% w/v) and chloramphenicol (100 µg.mL^−1^). Exconjugants were validated by PCR and inoculated in 5 mL M9 media + 2% (w/v) rhamnose for 1 hour at 37°C to induce toxin expression, followed by plating serial dilutions on M9 agar plates supplemented with 2% rhamnose and incubating at 37°C overnight. Successful double crossovers were confirmed by patching resulting colonies on LB and LB + chloramphenicol (100 µg.mL^−1^) plates. Chloramphenicol-sensitive colonies were screened by PCR and subjected to sanger sequencing to confirm deletion of target genes.

The *cpxA24* strain containing a stable in-frame deletion of *cpxA* residues 93-124 in the type strain ATCC 13047 was generated as above using the pTOX6 allelic exchange system (62). Regions of approximately 500 bp flanking the chromosomal deletion site were amplified using KOD and inserted into pTOX6. Conjugation and crossover selection was carried out as described above.

To complement the *cpxRA* deletion strain, *cpxRA* and their natural promoter were cloned and integrated at a single copy at the Tn7 site downstream of *glmS*. The CpxRA and the upstream intergenic region containing the promoter were cloned into the PmeI-AgeI sites of pTraCam-EcloGlmS plasmid and conjugated into Δ*cpxRA E. cloacae* as above. CamR exconjugants were verified by PCR. A Δ*cpxRA E. cloacae* strain expressing mVenus from the same locus was used as a negative control. The presence of the endogenous pECL_A and pECL_B plasmids was confirmed in all strains used for experiments.

### Bacterial growth curves

Overnight cultures of the *E. cloacae* wild type, Δ*cpxRA,* Δ*cpxRA*+CpxRA, Δ*cpxRA*+Ven or *E. cloacae cpxA24* were diluted to ODA_00_ = 0.05 into 25 ml LB plus the appropriate antibiotics. Cultures were incubated at 37°C with shaking, and absorbance readings were taken hourly for 8 h.

### RNA purification and sequencing

RNA was purified from exponentially growing cultures (OD_600_ = 0.5) of *E. cloacae* in LB alongside those of GAPEc grown to a similar OD_600_ and then left untreated or treated with 8 mg.L^−1^ gentamicin for 2 h. Experiments were performed on three separate occasions to collect biologically independent samples and purified together using Qiagen RNeasy kit following manufacturer’s protocol. RNA sequencing was performed at the Advanced Sequencing facility at the Francis Crick Institute. Samples were normalised to 500 ng and ribosomal RNA was depleted with RiboMinus Bacteria 2.0 Transcriptome Isolation Kit (Invitrogen, Cat no. A47335), followed by library preparation with KAPA RNA HyperPrep kit (Roche, kit code KK8541), according to manufacturer instructions. The adapter concentration was 7 µM and the libraries were amplified by 9 cycles of PCR. The quality and fragment size distributions of the purified libraries was assessed by a 4200 TapeStation Instrument (Agilent Technologies). Libraries were pooled and sequenced on the Illumina NovaSeq6000 in Single read 100 configuration to a read depth of at least 22 M.

### Identification of differentially expressed genes and pathway enrichment

Sequencing analyses followed established pipelines (65). Quality control was performed using fastqc (66) and read filtering with fastp (67). Briefly, reads were mapped to the *E. cloacae* ATCC 13047 reference genome (GCA_000025565, genome assembly ASM2556v1 from Ensembl using the GTF annotation file ASM2556v1.49c) using kallisto (v 0.46.2) in conda run on a Windows PC with a Linux subsystem (68), followed by tximport (v 1.26.1) (69) to generate length scaled TPM counts mapped to genes in *E. cloacae* in R/Bioconductor (v 4.0 or higher). Genes with counts >1 in all 9 samples were retained, and normalised prior to obtaining log2 cpm in edgeR (v 3.40.1) (70). The study design included 3 conditions (non-GAPEc, GAPEc & GAPEC+ gentamicin) and three repeats (9 total samples) with experiment as a blocking factor using “duplicateCorrelation” function in limma. Differentially expressed genes (GAPEc vs non-GAPEc, and GAPEc + gentamicin vs GAPEc contrasts) with false-discovery rate (FDR)-adjusted *P* value ≤ 0.01 with log2 fold-change ≥ 2 or ≤ −2 were generated with topTable in limma (v 3.54.0) (71).

gProfiler (online and R; (72)) was used to enrichment of pathways using GMT files obtained from Ensembl or a manually curated GMT file containing lists of transcription factors and their target genes (26). Transcription factors and their targets from the curated *E. coli* MG1655 genome in BioCyc (73) were mapped to the genome of *E. cloacae* ATCC 13047, and converted into a GMT file to use with gProfiler, including newly identified CpxRA targets in *E. coli* (26). Wrangling of these tables was performed using power queries in Excel. Gene Ontology (GO) terms and transcription factors with FDR-adjust *P* value ≤ 0.05 are shown in figures (74).

### Data collection and statistical analysis

For all experiments, a single mean was calculated from technical replicates (e.g., duplicate dilutions for MICs or serial dilutions for enumeration of viable CFU). Experiments were repeated independently on different days to obtain statistically independent means for different groups that were then compared in R (v 4.0 or higher) or GraphPad Prism (version 8.0 or higher). In the uncommon case of obtaining different MICs between technical duplicates, the highest concentration of the two was noted. Due to the unstable and heterogenous nature of heteroresistance, each colony of GAPEc and non-GAPEc on an agar plate were considered biologically independent, and typically 2-3 colonies were tested on a given day. Experiments were repeated independently on different days with different batches of GAPEc and non-GAPEc throughout this study, and all data were pooled for statistical analyses.

Student’s two-tailed *t* tests were used when comparing two groups. Non-parametric Mann Whitney test was used when comparing MIC of two groups, followed by false discovery rate (FDR)-based correction of *P* values for multiple comparisons (α = 0.05). Aligned rank transformation tool (ARTool package in R) was used for factorial ANOVA of MIC data without missing values and when F values of ANOVAs on aligned responses not of interest were 0. For more than two groups, factorial ANOVA (linear mixed effects models) was calculated with R packages lme4 (75) and emmeans (76) as implemented in grafify package (77). Mixed effects models used “Experiment” as blocking factor with random intercepts, and FDR adjustment (α = 0.05) was used to correct *P* values for multiple comparisons. For survival assays, log-transformations were typically required to ensure model residuals were approximately normally distributed following model diagnostics (e.g. using ggResidpanel (78)). Mean ± SD or median ± interquartile range (boxes indicating interquartile range (IQR), line at median, whiskers 1.5x IQR) are shown as indicated in figure legends. Data plotting used grafify and ggplot2 packages in R (79).

## Data and resource availability

The raw mRNA sequencing data are in BioProject PRJNA988129 and processed data under accession GSE236124 in the GEO database. Request for strains or plasmids generated in this study should be made to and will be fulfilled by ARS.

---

**Figure S1.**
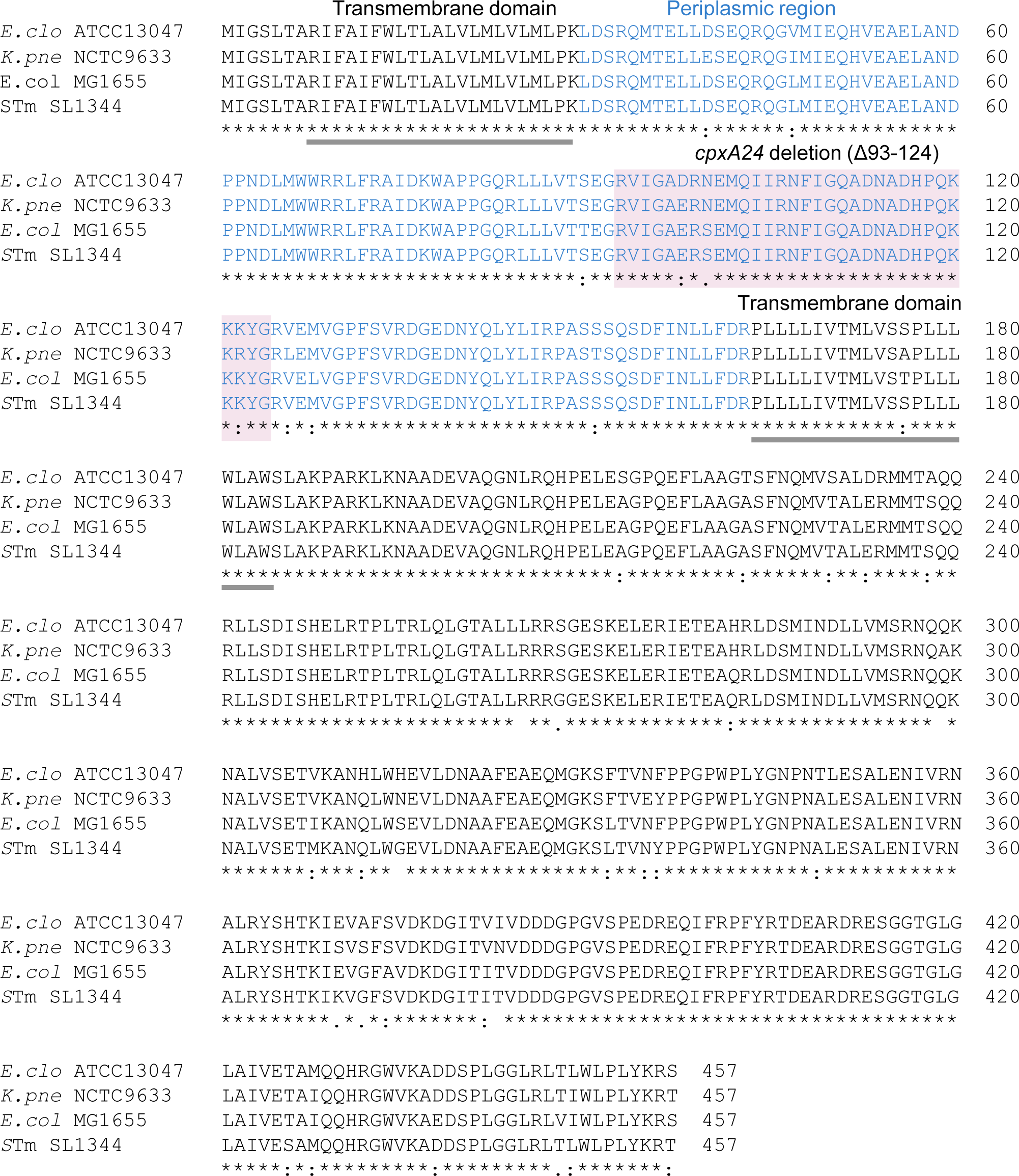
Sequence alignments of representative CpxA proteins. Amino acid sequence alignments of the CpxA proteins from *E. cloacae* ATCC13047 (*E.clo*), *K. pneumoniae* NCTC9633 (*K.pne*), *E. coli* MG1655 (*E.col*) and *Salmonella* Typhimurium SL1344 (*S*Tm). The *cpxA24* deletion site (aa 93-124) is highlighted (purple), along with the periplasmic region (blue) and transmembrane domains (underlined). The multiple sequence alignment was generated using Clustal Omega.

## Notes

### Competing Interest Statement

The authors have declared no competing interest.

### Summary of Updates

Updated incorrect author details and an author whose details were not saved in first submission.

